# To unscramble an egg: admixed captive breeding populations can be rescued using local ancestry information

**DOI:** 10.1101/2023.07.27.550812

**Authors:** Daniel J. Lawson, Jo Howard-McCombe, Mark Beaumont, Helen Senn

## Abstract

This paper asks the question: can genomic information recover a species that is already on the pathway to extinction due to genetic swamping from a related and more numerous population? We show that whole genome sequencing can be used to identify and remove hybrid segments of DNA, when used as part of the breeding policy in a captive breeding program. The proposed policy uses a generalised measure of kinship or heterozygosity accounting for local ancestry, that is, whether a specific genetic location was inherited from from the target of conservation. We then show that optimising these measures would minimise undesired ancestry whilst also controlling undesired kinship or heterozygosity respectively, in a simulated breeding population. The process is applied to real data representing the hybridized Scottish wildcat breeding population, with the result that it should be possible to breed out the domestic cat ancestry. The ability to reverse introgression is a powerful new tool brought about from both sequencing and computational advances in ancestry estimation. Since it works best when applied early in the process, important decisions need to be made about which genetically distinct populations should benefit from it and which should be left to reform into a single population.

## 1 Introduction

Recent advances in genomic technology have made high-density genotyping and sequencing conceptually accessible to conservation-led breeding programs. However, the benefits of doing so have not been clear because the majority of effort has been put into gaining value from more affordable technologies, primarily maintaining Pedigrees [1, 2, 3], and using low-density genetic markers [4]. Hybridization is a natural evolutionary force and the benefits of minimising it at varied species delineation levels are debated [5]. This is because species that can hybridize can be considered as populations of a higher-order species-complex, and mixing populations that were once structured may beneficially increase genetic diversity, without negative consequences for populations in the *future*. However, reducing anthropogenic hybridization (caused by domestic species or human-mediated introduction of wild species) is easier to advocate [6] as hybridisation is more likely to lead to problems. Genetic swamping of a threatened species by a domestic or human-associated species can cause inbreeding depression (low fitness due to decreased population size from wasted reproductive effort with the populous species) or outbreeding depression (low fitness of hybrids). This is especially important if the threatened species has an ecologically functional ecosystem role their more common breed do not perform.

In this article we do not assume that it is appropriate to reverse hybridization in any particular case - we simply propose a mechanism for doing so. Reversing hybridization is conceptually difficult as the genetic pieces of the two species become inseparable: you can’t ‘unscramble the egg’ [5], at least when using sparse genetic markers. We demonstrate here that high-density markers can achieve this, using breeding alone, by inferring *local ancestry*, i.e. assigning a particular population source to genetic segments. Observing local ancestry provides a revolutionary ability to breed-out past introgressed genomic segments. We propose ancestry-aware genomic analogue of the traditional kinship coefficient, which is the probability that both copies of a randomly chosen allele (i.e. variant of a genomic feature that varies between individuals) are Identical By Descent (IBD), here from a recent founder of the breeding population. This can be directly used in place of the pedigree in breeding selection [7].

Ancestry inference using software such as STRUCTURE [8] and ADMIXTURE [9] is common in conservation [10] using low density genetic markers such as Single Nucleotide Polymorphism (SNP) arrays or microsatellites [11], and targeted sequencing [12] such as ddRAD [13]. However, high-density markers reveal the details of Linkage Disequilibrium (LD) which allows statistical phasing [14] to estimate DNA inherited from each parent. Consequently shared haplotypes can be identified, that is DNA shared by descent between multiple individuals from a recent shared ancestor. Local ancestry specific to segments of each haplotype [15, 16] can then be inferred, for which we use the software MOSAIC [17], allowing detailed inference of population structure and identification of regions with anomalous ancestry indicating selection, e.g. in honeybees [18] and wildcats [19, 20]. These tools are important for understanding population structure [21] but so far have had little impact in conservation.

Captive breeding programs typically operate with smaller populations than would be ideal to maintain long-term population viability [22, 23]. The retention of genetic diversity is therefore the focus, targeting an effective population size (*N*_*e*_) of at least 500 [24], a threshold used by the Convention on Biological Diversity (CBD) [25] to describe a healthy population. Lack of diversity leads to disproportionate heritable disease burden, for example inbreeding depression in Italian wildcats [26]. For this reason, breeding management — as exemplified by the popular PMx software [11] for decision support, as well as Vortex [27] for population viability analysis — emphasise minimising kinship. Kinship is the proportion of the genome that is identical due to recent shared relatives between parents of an individual [28]. Kinship is not the only way to measure inbreeding — alternatives include using (sparse) molecular markers to measure Allele Sharing [13], or the heterozygosity [29, 10], which is the proportion of markers that are different. Because these sparse genomic measures incompletely characterise variation, the benefits of quantifying them are debated.

Endangered species that experience anthropogenic hybridisation with more numerous species, typically domestic or human-associated, are under threat of extinction via genetic swamping [30, 31], i.e. the replacement of their genome by that of a more numerous related species or population. Partridges and wolves [32] experience this problem, and it is a threat to captive breeding programs including African penguins [33], crocodiles [34], and in our case, the Scottish wildcat. Such species pose a particular challenge for conservation because the breeding program may not have been in place before hybridization. This is the case for the Scottish wildcat, for which breeding cats possess an average of 18% admixture from domestic cats in the captive population, and higher in the remaining wild population [35]. There are no unadmixed individuals, but we have recently shown [19] that a near-complete genome exists across the population. The current protocol excludes individuals with more than 25% domestic ancestry [36] based on measuring 35 well-chosen genomic markers. Yet we will show that if it is deemed desirable to do so, hybridization of this magnitude and timescale can be recovered to close to pre-hybridization levels by inferring local ancestry for dense markers.

The challenge for conservation in such cases is that there is currently no tool to remove hybridization. Controlling kinship alone will not remove the introgressed genome. Trying to maximise genetic diversity using molecular estimators could paradoxically make things worse due to the introgressing species being genetically distinct from the target species, so that selection for diversity increases the frequency of introgressed alleles. Few measures suitable for breeding choices whilst accounting for local ancestry are available. Much prior work focuses on the estimation of heterozygosity of local ancestry itself, e.g. [37, 38]. Heterozygosity of F2 hybrids has be used to compute hybrid incompatibility [39], and runs of homozygosity have been decomposed by local ancestry [40]. Ancestry-specific *F*_*ST*_ has been linked to kinship [41] for the purpose of estimating population structure. In contrast to previous work, we will introduce a local ancestry ‘mask’ for computing ancestry-specific heterozygosity or kinship in admixed individuals.

Figure 1 illustrates how the approach works. We observe that recent tools for local ancestry estimation could be used for a breeding program. It is relatively straightforward to estimate ancestry from wholegenome data in the well-diverged species of conservation interest (if a reference alignment and suitable proxy populations are available), so we can assume that the ancestry status of haplotypes are known (demonstrated for wildcats in Figure S1). We therefore use simulation to examine how this can be used in a breeding program. We examine different choices of selection measure for the breeding program, and consider the genetic diversity consequences for the species. Existing breeding-decision rules for kinship or heterozygosity can be extended to use this local ancestry information by ‘masking’ out ancestry from the undesired species, which reduce to familiar measures in the absence of introgression. We find that correct management using our proposed local ancestry measures can turn around the introgression status of a population in many circumstances.

**Figure 1.**
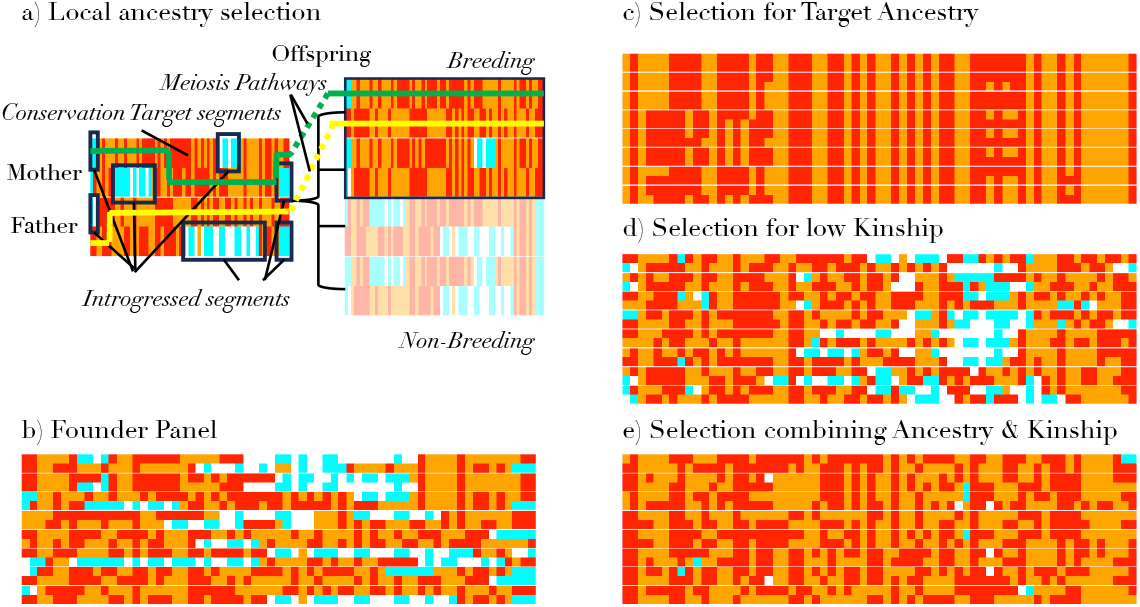
Concepts in this paper. a) The process of meiosis takes the two haplotypes from each parent and makes a single random haplotype; the yellow and green lines are one possible realisation. The conserved target ancestry (orange, red with mutation) can be separated from Introgressed ancestry (white, cyan with mutation). This allows selection of the most suitable offspring for the next generation, who each receive different random haplotypes. b) Initially we start with a panel of relatively diverse and introgressed individuals. c) If we select against introgression, we lose diversity in the genome. d) If we select to minimise kinship, we fail to remove introgression. e) If we select for diversity with knowledge of local ancestry as in a), we can balance both pressures.

## 2 Results

### 2.1 A simulated breeding program

The experimental design considers a hybridized captive population with genetic components from two original sources, the ‘target’ population for conservation and the ‘introgressing’ population. We simulate a founder captive breeding population (Section 4.9) of *N* = 150 diploid individuals at *L* = 500 linked genetic positions (called loci) over 5 Morgans (i.e. 5 crossovers are expected per generation per parent) of genome in which two generations of admixture occurred 10 generations before the captive population was formed. There are two generations in the establishment of the captive population, the first being a bottleneck due to sampling wild individuals, and the second being a selection of the least introgressed captive individuals. We then simulate the captive population forwards in time, making breeding decisions based on the ‘ranked mean’ selection procedure (Section 4.10), which has been shown to be appropriate for kinship (the ‘ranked MK’ procedure) for discrete populations [11]. Briefly, this procedure iteratively ranks individuals from ‘best’ to ‘worst’ in terms of optimising pairwise breeding measures in the next generation. It then assigns a random number of offspring, from the best pairs downwards, until the program reaches capacity in the next generation (here *N* = 150).

Firstly we consider admixture unaware breeding measures, which contain information that can be acquired without local ancestry information. As a baseline, we consider choosing individuals randomly (*R*). This is compared to breeding selection measures which optimise an expected score in the offspring, conditional on the parents’ genome. We consider minimising the expected kinship (*K*; Section 4.1), which we compute not from pedigrees (which we call theoretical kinship) but conditional on the local genome (which we denote empirical kinship), specifically by asking if the genome segment derives from the same founder haplotype. The expectation here is over the values of kinship that offspring with each other individual might have, and due to the ranked MK process is taken only over ‘better’ individuals. We also maximise expected heterozygosity (*H*; Section 4.2). We contrast this with minimising expected introgressed admixture (*Q*; Section 4.3).

The same measures that are used as a performance measure can also be used as a score to quantify performance in the realised population. When used as a score, these are denoted as *S*^*Q*^ for admixture, *S*^*H*^ for heterozygosity, and *S*^*K*^ for kinship. Scores are distinct from expected measures because they are observed in the current population, not an average over possible pairs. As shown in Figure 2, no measure is satisfactory: minimising introgressed admixture leads to a diversity collapse (both for heterozygosity and kinship). Conversely, minimizing kinship best maintains the population as it was, whilst maximising heterozygosity increases introgression as the introgressing population is diverged from the target.

**Figure 2.**
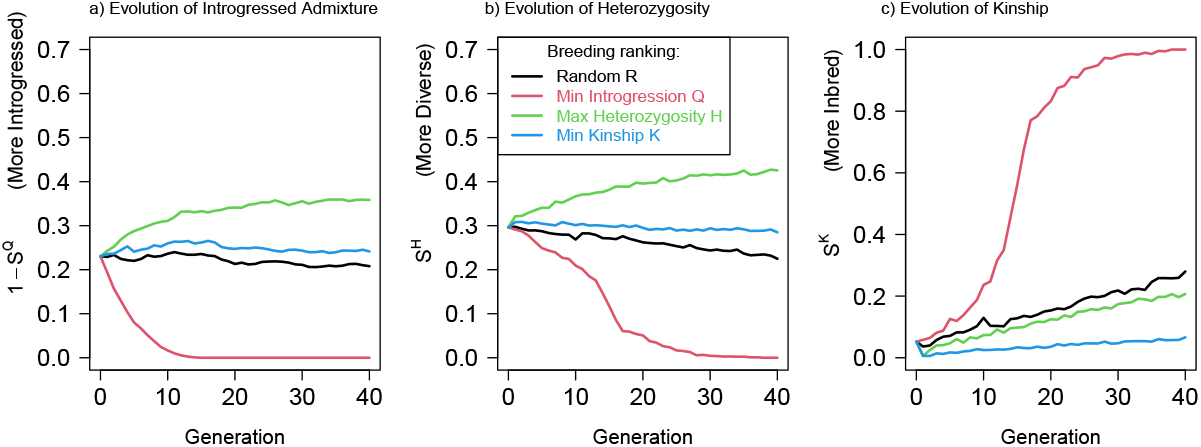
Simple measures cannot simultaneously remove introgressed ancestry and maintain diversity. Here we either breed random pairs (R), attempt to minimise kinship (K) or introgression (Q), or to maximise heterozygosity (H). We quantify their performance over time with scores for a) introgressed admixture fraction 1 *− S*^*Q*^, b) heterozygosity *S*^*H*^, and c) kinship *S*^*K*^.

Local ancestry information allows us to assign ancestry to specific genomic loci. In order to balance population ancestry and diversity, we introduce population heterozygosity (PH, Section 4.4) and population kinship (PK, Section 4.5). These measures exploit being able to assign specific position on a genome to a population and can be interpreted as the mean value (of *H* or *K*) at loci on haplotypes where *both* copies were inherited from the target population. Introgressed loci are ‘masked out’, that is, treated as heterozygosity 0 and kinship 1. When used for prediction of the next generation, we weigh loci by the probability they take each ancestry. Population heterozygosity and population kinship are easily evaluated as scores *S*^*PH*^ and *S*^*PK*^ respectively, averaged across realised individuals.

The use of population measures punish hypothetical offspring with one or two introgressed haplotypes equally. To address this, we also consider weighted population heterozygosity (WPH) in Section 4.6 and weighted population kinship (WPK) in Section 4.7), in which a weight *δ* is placed on the population measure and 1*−δ* is placed on expected admixture *Q*. Throughout we use *δ* = 0.5 for heterozygosity, with the appealing interpretation of placing relative weight 0 on two introgressed alleles, 1 on a single target population allele, 2 on homozygous target population alleles and 4 on heterozygous target population alleles.

The result of using each of these local ancestry measures are shown in Figure 3. All suggested local ancestry measures control introgression nearly as well as minimising it directly. Simultaneously, they maintain heterozygosity and kinship adequately overall, and strongly improve them for genetic material from the conservation target population. Overall, weighted and unweighted population kinship measures perform similarly, and are desirable if kinship is to be minimised. Weighted population heterozygosity could be argued as preferable as it maintains heterozygosity above the starting value. Importantly, and perhaps surprisingly, if we allow the realities of the breeding program to allow only 60% of breeding recommendations to be followed (60% ‘compliance’; Section 4.10.2), with other parents chosen randomly, the recovery is still dramatic with strong control over admixture *Q* and no severe consequences for heterozygosity or kinship.

**Figure 3.**
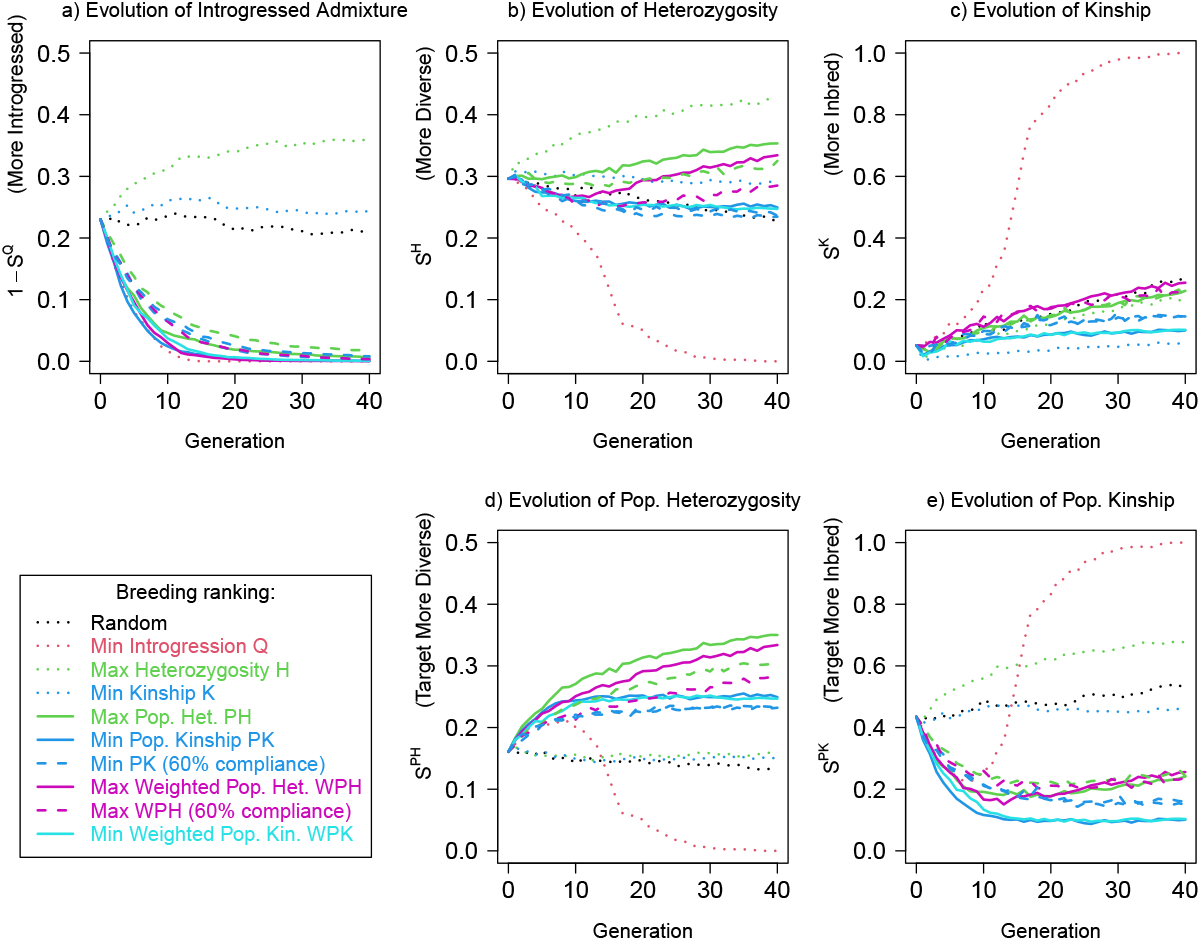
Local ancestry-based measures can simultaneously remove introgressed ancestry and maintain diversity. The curves from Figure 2 are shown as dotted curves, and we add curves attempting to minimise population kinship, population heterozygosity, and their weighted versions. We also add ‘60% compliance’ curves for which the recommendation is followed 60% of the time, and a random choice of parent is made otherwise. We quantify performance over time with scores for a) introgressed admixture fraction 1 *− S*^*Q*^, b) heterozygosity *S*^*H*^, c) kinship *S*^*K*^, d) population heterozygosity *S*^*P H*^, e) population kinship *S*^*P K*^.

Having established that the genomic measures work in principle, we now more thoroughly consider the details of the simulation scenario, which can affect the speed that the breeding program operates and the final outcome in terms of diversity. We focus on the best two options, selecting for population kinship (PK) or weighted population heterozygosity (WPH). Figure 4a,e) show that heterozygosity decreases with initial introgression in proportion to the diversity of target ancestry present in the original panel. Kinship likewise increases due to fewer founders being available at each locus. The population-specific scores closely match the introgression-free ones, and only deviate when there is limited power to remove introgression over the timeframe (here 20 generations). Both selection options are adequate, with heterozygosity selection increasing both kinship and heterozygosity slightly.

**Figure 4.**
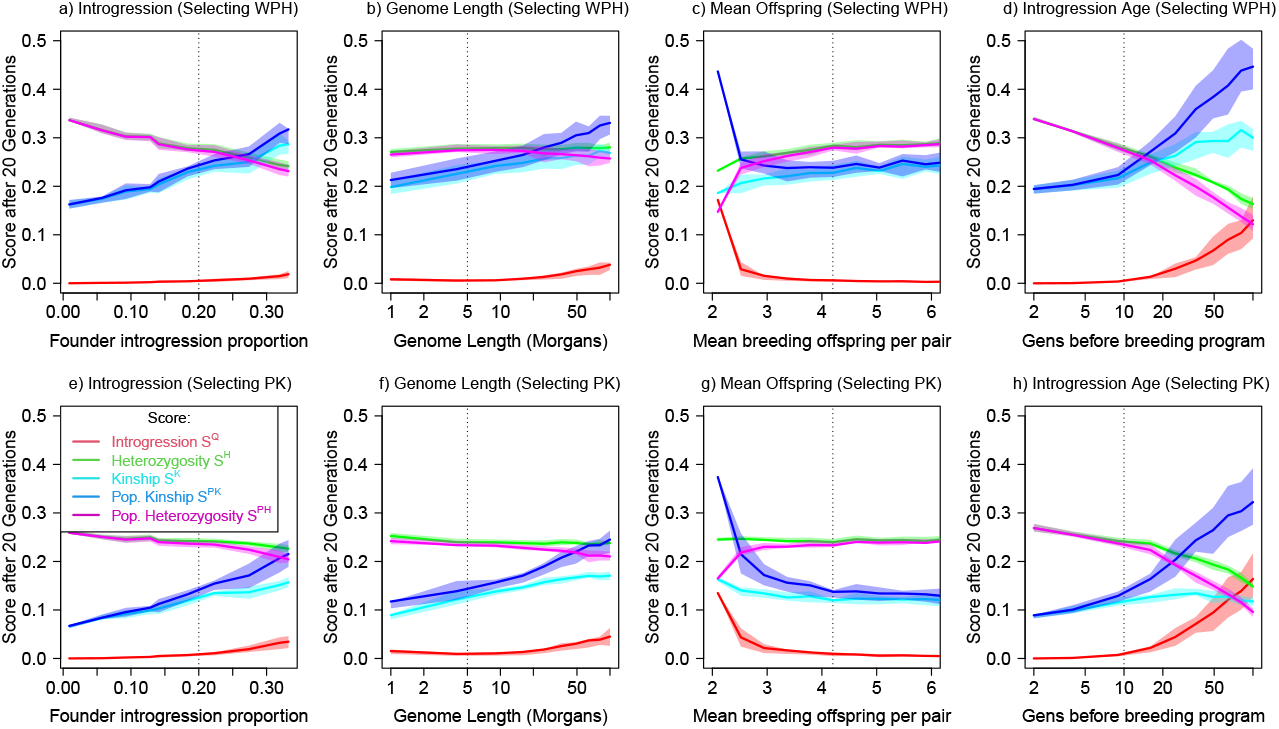
Impact of the simulation parameters on the scores after 20 generations of selective breeding using (top) the weighted population heterogeneity (WPH) score measure and (bottom) population kinship (PK). This is in populations with otherwise the same demography as Figure 2-3 (whose parameters are shown as vertical line). Here we consider changing a,e) the initial amount of introgression observed in the founder captive population, b,f) the total quantity of genome in Morgans, retaining the same number of SNPs, c,g) the average number of offspring who survive to adulthood and are available for the next generation of breeding program, d,h) the number of generations between admixture and the creation of the breeding program. The scores are the 5 shown as separate panels in Figure 3, colour-coded to their corresponding breeding measures.

Figure 4b,f) show that increasing the genome length decreases the selection efficiency, with a corresponding increase in introgression *Q*. Kinship, heterozygosity and population heterozygosity scores are not strongly affected. Population kinship is increased at longer genomes. We hypothesise that this is due to having more introgressed segments remaining, and a decreased diversity of candidates as time progresses, meaning that replacing introgressed segments later comes at a higher kinship cost. Animal conservation target species are likely to have less than 50M (Morgans) of genome, with sex-averaged genetic map lengths being surprisingly consistent across genera; for example, the mouse (16.3M [42]), cat (43.7M), superb fairy-wren (16.2M [43]) and stickleback (12.5M [42]) genomes are all on the same order of recombination length to humans (35M [44]), and in all cases genome length is not a barrier to removing introgression after 20 generations. Larger brood sizes typically lead to more options for breeding and therefore a more efficient breeding program. Figure 4c,g) shows that the number of offspring who themselves survive to breeding age should be above around 3.

One of the more dramatic changes to a scenario is the number of generations before admixture. Figure 4d,h) highlights how vital it is to make an early intervention: after about 10 generations, the ability to reduce introgression decreases, and the resulting cost in kinship is increasingly high. This is because the introgressed DNA exists in increasingly small segments as the introgression event fades into the past. We emphasise that the initial introgression level was controlled to 20%, and that we are examining the population after 20 generations of the breeding program. Breeding for longer can to some degree offset a delay.

### 2.2 Application to the Scottish Wildcat

Here we apply the breeding simulation to the best approximation of the Scottish wildcat breeding population for which data is available. We emphasise that these data are considerably more introgressed than the breeding program individuals, due to containing a high number of wild-living individuals. The following information is given in more detail where the data were first presented [20, 19]. The Scottish wildcat is the north-westernmost population of the European wildcat *Felis silvestris* which is distributed in separated populations across Europe. It is related to the domestic cat, which evolved from *Felis lybica* which is distributed across north Africa and the middle east. Domestic and wildcat species are genetically quite distinct having diverged around 200k years ago [45], with *F*_*ST*_ = 0.46 and therefore the two species are quite separable. In Britain, the species were reproductively isolated until the mid 1900s, even though both species were present since the introduction of the domestic cat during the Roman occupation.

We start with a dataset of 36 cats [19] and selected the 24 genomes with the highest Scottish wildcat genome fraction. We applied realistic choices for parameters (number of lifetime offspring per pair; carrying capacity) matched to the breeding program (Section 4.11). In brief, this dataset was generated from a combination of 6 captive cats in the Scottish wildcat breeding program, plus 30 wild living cats collected as part of a diversity panel. Introgression was confirmed in the captive population using ancient DNA as well as historical museum samples. The captive cats had an average domestic cat introgression of 0.18, whereas the wild-living cats ranged from 0.11-0.82.

To replicate a genomic panel that we might find in the captive population, we annotated each of these individuals as founders, selected the top 24 by genome-wide wildcat genome proportion (maximum 0.53, mean 0.18) and performed a single breeding step to create a ‘captive population’ of *N* = 150 individuals. The whole cat genome is 43.7 Morgans [46], and we analyse chromosome E3 which consists of 1.26 Morgans observed over 17720 SNPs. Because the introgression is relatively recent, the chromosome varies considerably from genome-wide admixture fractions, with a mean wildcat ancestry per haplotype of 0.60 and a range of 0.14-1.00.

Figure 5 shows the results of the same breeding process described for the simulated data for the case of Scottish wildcat introgression. Parameters have been matched to the breeding program where possible: the mean number of offspring per breeding pair is 4.2 which is the lifetime expected number per female as observed in the stud book [47] and 150 individuals is the breeding program size. Qualitatively the results agree with results on the simulated haplotypes, with the local-ancestry measures successfully controlling introgression and maintaining diversity. Due to the lower number of founders and the higher initial introgression, there is a separation between the scores, with WPH removing introgression significantly faster than selection for the PK kinship measure (especially over a 5-10 generation horizon) but at the price of increased kinship. In this case the ability to comply with breeding recommendations does start to have an impact; an 80% compliance rate in qualitatively similar to full compliance and can achieve eventual wildcat ancestry recovery, whereas a 60% compliance rate leads to considerably slower recovery.

**Figure 5.**
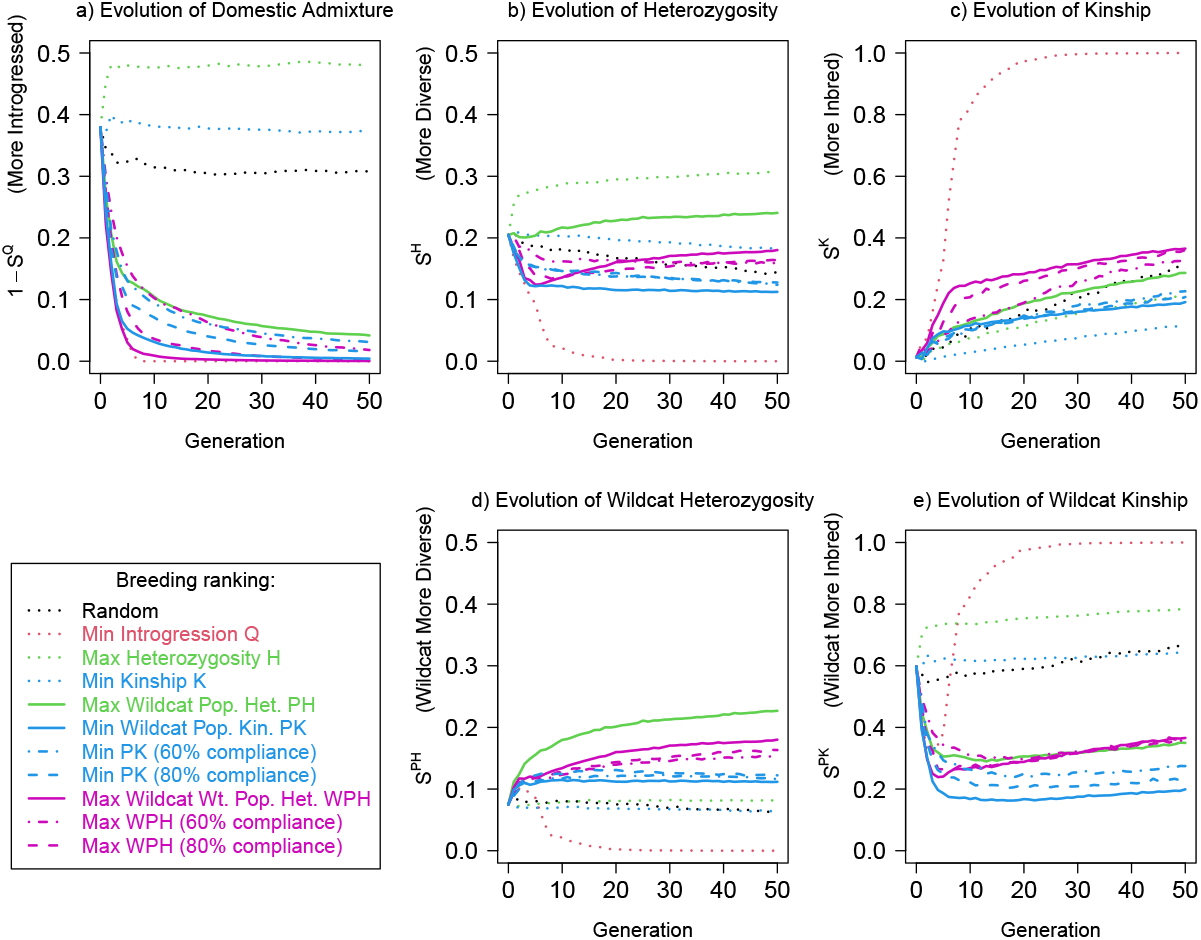
Results for a putative Scottish wildcat breeding program based on real genetic data and local admixture estimation for 28 founders and a captive population capacity of 150. Curves are as Figure 3, showing traditional measures as dotted lines and best-case local ancestry measures as solid lines. ‘Real world’ limited compliance (80% dashed, 60% dot-dashed) with breeding recommendations are also shown. The scores are a) domestic cat admixture fraction 1 *− S*^*Q*^ (note that these founders have higher introgression that the captive breeding program); b) heterozygosity *S*^*H*^, c) kinship *S*^*K*^, d) wildcat population heterozygosity *S*^*P H*^, e) wildcat population kinship *S*^*P K*^.

## 3 Discussion

We have shown that traditional measures used for breeding decisions in captive breeding programs are inadequate for populations that experienced introgression prior to the initialisation of the captive population. For such populations we must be very careful not to mistake introgression for diversity. However, a variety of measures based on using ‘local ancestry’ to determine the introgression status of each genomic region are extremely effective at reducing introgression whilst simultaneously maintaining genetic diversity, whether measured in terms of kinship or heterozygosity.

For the case of the Scottish wildcat, we have shown that the additional information required, i.e. local ancestry information, is quite straightforward to access in practice [19] for species for which a reference alignment and recombination map are available. From relatively low-coverage whole genome sequencing data we would be able to accurately infer haplotypes via statistical phasing, and attribute local ancestry via chromosome painting. As a pilot demonstration, we were able to perform the breeding program *insilico*, not from the active captive population (for which sequencing is unavailable) but using a diversity panel. Simply by sequencing the active breeding population of a program, we could apply our pipeline to improve breeding pair selection. The complexity for a breeder over current practice of using any such measure, for example population kinship instead of kinship, is low and the gain in terms of purging introgressed genetic segments is dramatic.

A critical component of a successful reversal of hybridization is timing. It is only possible to undo recent hybridization, as the required duration of the breeding program will grow in proportion to the delay, which places strong practical and cost constraints on the process. ‘Unscrambling the egg’ of hybridization takes as long as it was left to scramble so there are considerable cost savings and practicality benefits in acting early, if the decision is to act at all. For the case of the Scottish wildcat diversity panel data, estimates for individuals range from between 4.7 and 17.9 generations before present, with a mean estimate of 8.6 (95% CI 8.3 - 9.8) generations. This scale of hybridization is consistent with a predicted potential recovery over 10-20 generations of breeding program, provided that compliance with the breeding recommendations is not too low. Without sequencing the captive cats, we can only say that the option is promising; the ‘future captive breeding population’ simulated here differs in both the initial admixture and diversity, the genome length simulated, and in the particulars of generation structure. These particulars are required to tell how stringently breeding recommendations must be followed to achieve adequate recovery in practice.

There are still however many challenges remaining. A primary one is funding. Whilst the cost of sequencing continues to drop, it remains non-negligible and needs to be maintained over time. A second challenge is to develop software that incorporates these genomic features whilst maintaining all of the small subtle details that allow the simulation to represent a breeding program, including the correct generational structure, the practicalities of creating mating pairs, and so on. A third is the ticking clock of diversity within species: as captive populations are maintained in small populations, kinship inevitably increases. The health consequences of kinship as measured this way are unclear - in our simulated breeding conditions we were able to maintain heterozygosity even in the face of rising kinship, which is present in increasingly small segments of Identity by Descent.

A final challenge is that we are becoming increasingly aware of genomic features of captive populations that require special treatment. We have not modelled inbreeding depression beyond a simple increased mortality in the first year. In modelling tools such as PMx [11] in which the genome is not available, recessive disorders are modelled as independent random events occurring with rate proportional to the kinship coefficient. We have the advantage of observing the kinship for specific loci, which might in principle be used to infer regions of increased and decreased inbreeding risk. Similarly, outbreeding deperession may not be uniform along the genome. Some genes may be advantageous to receive via introgression, most notably immune genes. For example, the major histocompatibility complex (MHC) region which regulates immune response, may have been selected for domestic cat ancestry in wildcats [19], due to the high domestic cat disease burden. Breeding programs may wish to take this into account. This is related to the desired outcome - how much introgression needs to be removed? The answer may well be related to real-world measures of both fitness and ecological function, which are yet to be robustly related to genomic properties. It is possible that widespread linkage of pedigrees with phenotypes and genomic quantification of populations over time can address these problems, but that is yet to be proven.

The potential that such fine-scale genomic information brings to conservation genetics implies that a new generation of tools may use far more detailed representation of the genome for selection than the simple scores defined here. Some caution may therefore be needed before the tools are mature, but the advantages of acting early to save the diversity that is present needs to be weighed against the risks.

## 4 Methods

Throughout we will follow at generation time *t* a genetic panel *G*(*t*) making up a 3D array of dimension *N* (*t*) *×* 2 *× L. G*_*ial*_(*t*) is indexed by the individual *i* = 1 … *N* (*t*), the haplotype copy *a* = 1, 2 and the genetic loci *l* = 1 … *L*, and takes value 0 for reference and 1 for alternate alleles. We will also follow an ancestry panel *A*(*t*) of *N* (*t*) *×* 2 *× L* local ancestry values, taking value 1 for our Target ancestry and 0 for Introgressed ancestry. To ease the notation, where possible we omit the time variable when referring to the current generation. We also keep a simple population size schedule where *N* (0) = *N*_*F*_ is the number of founders and the captive population size *N* (*t*) = *N* is fixed for *t >* 0.

*G*(*t*) and *A*(*t*) are tracked directly from one generation to the next using the process of meiosis (see Section 4.8).

### 4.1 Kinship Coefficient and local pedigree

For the simulation study, we will follow the ‘local pedigree’ or founder panel *F* (*t*) of *N* (*t*) *×* 2 *× L* founder sources, describing the founder haplotype of each genomic locus *l* from a founder population forwards in time. For individual *i, F* is initialised to *F*_*ial*_(0) = 2*i − I*(*a* = 1) with *a ∈* (1, 2), i.e. the whole of each haplotype in the founder population is assigned a unique value from 1 to 2*N*_*F*_ . *I*(*x*) is an indicator function, which for binary *x* takes value 0 when *x* is false and 1 when *x* true. Like *G* and *A, F* is tracked during meiosis (Section 4.8).

By tracking the founder panel we can compute the (empirical) expected kinship coefficient *K*_*ij*_ between individuals *i* and *j*. The kinship coefficient is the probability that two randomly chosen alleles are Identical By Descent (IBD) and is typically conditioned on the pedigree alone (which we call the theoretical kinship). However, as we observe the genome it is useful to condition on it (which we call the empirical kinship or simply the kinship). The expectation is taken over potential offspring and is the average over each specific locus *l*. The *expected kinship coefficient* 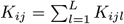 is to be written in terms of *F*_*ial*_, the founder ancestor of haplotype *a* from individual *i* at locus *l*:

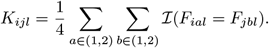

Although *F*_*ial*_ is not directly estimable from data, the IBD status *I*(*F*_*ial*_ = *F*_*jbl*_) is [7, 13], so *K* can still be estimated at additional computational cost.

The expected kinship coefficient *K*_*ij*_ is used for decision making. The score that we report for performance is the empirical mean for realised individuals, i.e. the ‘kinship score’ is 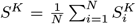 where the individual kinship is:

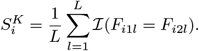

Kinship is to be minimised. Throughout we define expected values for pairing for decisions as matrices using capital letters and scores based on them as superscripts on *S*, i.e. *K* and *S*^*K*^.

### 4.2 Heterozygosity

Analogously to kinship, we can define heterozygosity as both a score and a pairwise expectation for the offspring of each pair of individuals. Given a pair of individuals *i* and *j*, the *expected heterozygosity* of their offspring is 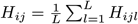 where for locus *l*:

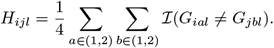

The heterozygosity score 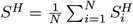 where 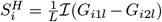, i.e. takes value 0 if the two copies are the same and 1 otherwise. Heterozygosity is to be maximised.

### 4.3 Admixture

It is natural to quantify ancestry via the proportion of ancestry from the target population, by convention called the ‘Q’ statistic. We have defined *A*_*ial*_ = 1 if the *a*-th haploptype from individual *i* at locus *l* is from our target population.

The *expected admixture* is 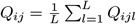 where

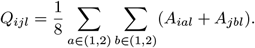

The admixture score is 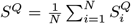 where 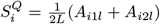 and is to be maximised. We will report in terms of the introgression score 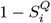 which is to be minimised.

### 4.4 Population Heterozygosity

We now condition on local ancestry to make a population-specific score. This is slightly more transparent for heterozygosity than for kinship, and for the score rather than the expected value in the next generation, though both follow analogously. Population heterozgosity (PH) is a product of two terms, whether both copies come from the target population, and whether the SNP is heterozygous conditional on this. We define the target ancestry diploidy *I*(*A*_1*l*_ + *A*_2*l*_ = 2) as an indicator function taking value 1 if both haplotypes are derived from the target population, or 0 otherwise.

The *expected population heterozygosity* 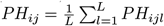 *i*s defined for a pair of individuals *i* and *j* and the expectation is taken over their potential offspring:

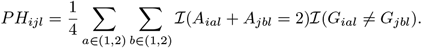

The PH score 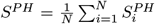 is the average over realised individuals’ scores 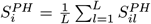 across the genome, where:

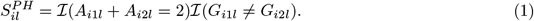

Population heterozygosity is to be maximised. It is equal to the heterozygosity if all copies of all loci are from the target population.

### 4.5 Population Kinship

The population kinship is constructed as the population heterozygosity, but more care is needed to ensure that the target population ancestry is selected to increase whilst kinship is selected to decrease. We wish to only reward loci that both are from the target ancestry and have different founders *ℐ*(*F*_*ial*_ ≠ *F*_*jbl*_). The *expected population kinship* 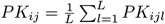 is therefore defined as:

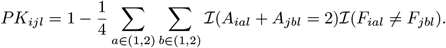

Similarly, the *population kinship score* 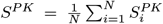 is the sum over the individual realised population scores 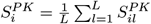 where the per-locus score is:

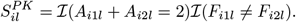

Population kinship is to be minimised and is equal to the kinship if all copies of all loci are from the target population.

### 4.6 Weighted Population Heterozygosity

There is no reason to expect that the best-performing rule should only consider ancestry homozygotes for the target population - particularly for rare SNPs, it might instead be better to reward any ancestry. We therefore consider the *weighted population heterozygosity* 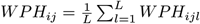 where:

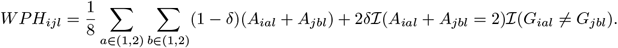

WPH is to be maximised. If we set *δ* = 0 this returns to the target population admixture *Q*_*ijl*_, and with *δ* = 1 this becomes the population heterozygosity. We use *δ* = 0.5 throughout, which rewards heterozygosity equally to ancestry dosage, and which leads to the per-locus value 0 for no target alleles, 0.25 for a single target allele, 0.5 for two identical target alleles, and 1 for heterozygous target variants.

### 4.7 Weighted Population Kinship

The same procedure can be applied to obtain the *weighted population kinship* 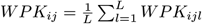 where:

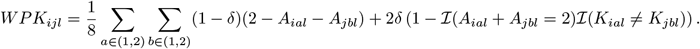

This transitions between selection against introgression and selection against kinship. WPK is to be minimised.

### 4.8 Simulated meiosis

Individual genomes are observed over a single chromosome with genome length *D* (measured in Morgans, default 5) at *L* single nucleotide polymorphisms (SNPs; default 2000). *D* is the average number of recombination events separating parent haplotypes from the child haplotype. Let *R*_*l*_ for *l* = 1, …, *L* denote the recombination position of the *l*-th SNP, i.e. taking values in the range (0, *D*). In the simulation we evenly-space SNPs so that SNP *l* has recombination position *R*_*l*_ = *lD/L*.

Meiosis begins by producing a gamete for each parent. Let *a*(*l* = 1, …, *L*) be a vector of length *L* taking value 1 or 2 to denote which of the two haplotypes is being inherited at position *l*. We generate *a* via an independent crossover recombination process:

1. Initialise:
  a. Set the initial position *p*(*b* = 1) = 0 where *b* indexes the recombination segments, starting at *b* = 1.
  b. Choose an initial haplotype *h*(*b* = 1) *∼ U* (1, 2), i.e. start with either the first or second haplotype with equal probability.
2. Until *p*(*b*) *> R*_*L*_, i.e. the whole chromosome is processed:
  a. Generate a segment inheritance length *l*(*b*) *∼* exp(1).
  b. Let *e*(*b*) = *p*(*b*) + *l*(*b*) be the end point of the inherited segment. For every SNP in the segment, i.e. for *l* such that *p*(*b*) *≤ R*_*l*_ *< e*(*b*), set *a*(*l*) = *h*(*b*).
  c. Set *h*(*b* + 1) = 3 *− h*(*b*), i.e. switch from haplotype 1 to 2 and vice-versa. Set the next start point to be the current end point, i.e. *p*(*b* + 1) = *e*(*b*), and move on to the next segment setting *b ← b* + 1.

We generate the gamete *a*_*i*_ for parent *i* and *a*_*j*_ for parent *j*, tracking *G, A* and *F* . Specifically for the *k*-th offspring in generation *t* + 1 with parents *i* and *j*, we set 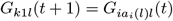, and 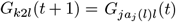 likewise for *A* and *F* .

### 4.9 Simulated population

The simulation uses two related populations *P*_*A*_ and *P*_*I*_, the target and introgressing populations respectively, who share ancestry via a common ancestor *P*_0_. The common ancestor has SNPs *l* = 1 … *L* taking value 0 by default or 1 with frequency *f*_*l*_ *∼ U* (0.05, 0.95) and the population frequencies follow the Balding-Nichols model [48] with Fst *F*_*A*_ and *F*_*I*_ respectively (both assumed 0.2):

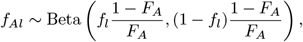

with expectation 𝔼 (*f*_*Al*_) = *f*_*l*_ and variance Var(*f*_*Al*_) = *F*_*A*_*f*_*l*_(1 *− f*_*l*_). The same model is applied to the introgressing population *P*_*I*_ . The genetic separation between populations *P*_*A*_ and *P*_*I*_ is by construction *F*_*ST*_ = *F*_*A*_ + *F*_*B*_ = 0.4.

We form an introgressed *wild* population as follows. First we generate a genetic panel *G*_*A*_ of 2*N*_*A*_ haplotypes to form the pre-introgression target population, matched with an ancestry panel *A*_*A*_ with every entry taking value 1. We also form an introgressing panel *G*_*I*_ of 2*N*_*I*_ haplotypes to form the introgressing population, matched with *A*_*I*_ with every entry taking value 0. We then impose a sequence of introgression rates *α*(1), …, *α*(*T*_*init*_), using *T*_*init*_ = 10 and *α*(1, 2, 3, …, 10) = (0.2, 0.2, 0, …, 0), i.e. introgression happened 10 generations in the past and lasted 2 generations. We then perform random mating with the specified introgression rate. Random mating uses the Wright-Fisher process, i.e. each child chooses two random parents, with probability *α*(*t*) uniformly from the introgressed panel and with probability 1 *− α*(*t*) from the target panel. The children are generated by the meiosis process described in Section 4.8.

We then perform two additional generations to form a more realistic *captive* population: a *bottleneck* to replicate a small number of founder individuals being sampled for introduction into the program, and *selection* for the most representative of those founders. In the bottleneck step, a fraction *p*_*bottleneck*_ (=0.25 here) of individuals are sampled for uniform random breeding. These individuals are indexed to define the founders. To form the most promising breeding population, in the captive step, a fraction *p*_*captive*_ (=0.25 here) are selected by minimum introgression proportion and finally bred (using the Wright-Fisher process) to form the *captive* population containing *N* individuals. Introgression therefore occurred *T*_*init*_ + 2 generations ago.

### 4.10 Simulated Breeding Program

Here we describe how captive individuals are chosen for breeding within the captive breeding program. A breeding program contains many details that will differ from a simulation. Our conclusions are robust to the particular design of the generational structure (with sensitivity analyses described below.) For ease of exposition we focus on a discrete-generation process in which a random number of offspring are produced per breeding pair *n*_*ij*_ ∼ *ℱ*_*n*_ = Poisson(*λ*) (with mean 1*/λ* = 4.2 here).

#### 4.10.1 Ranked Breeding Rule

Breeding programs can use a range of schemes to pair individuals [11] based on minimising (or maximising) a breeding score *B*_*i*_. The breeding score should not be considered as a property of an individual, but is instead based on a pairwise matrix *B*_*ij*_ that can be a function of the next generation being constructed. The expected (i.e. mean) kinship 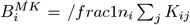 is a popular choice, where the expectation is taken over *n*_*i*_ individuals determined by a selection process (see below). The expected admixture *Q*_*i*_, inbreeding coefficient, identity-by-state, identity-by-descent, or similar can all be used. [11] describe processes based on ranking, with elimination of pairs that are too genetically similar. For discrete generations they found that ‘ranked selection’ was appropriate. We implement their scheme without sex constraints for simplicity. This creates a ‘breeding score’ *B*_*i*_ for each individual, that is updated to reflect the other potential parents expected to be in the next generation. This is a good approach because the breeding scores computed can be seen as expected values over the entire captive population and hence reflect the ‘value’ of the individual to the whole breeding program.

The procedure is as follows (assuming we are maximising *B*):

1. Compute *B*_*ij*_ for the current generation.
2. Set the parent stack *s* = () empty, and the remaining list *r* = (1, …, *N*) containing all individuals.
3. Iteratively construct the parent stack, from least promising to most promising. For *t* in 1, …, *N* :
  a. Compute the breeding score 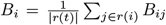 which is an average breeding score over the |*r*(*t*)| remaining individuals;
  b. Select the worst individual *k* = argmin_*i∈r*_*B*_*i*_;
  c. Add *k* to the parent stack, i.e. set *s* = (*s, k*);
  d. Remove *k* from the remaining list, i.e. set *r* = (*r \ k*).
4. Perform pairing from most promising to least promising. Set pair number *t* = 1, current offspring count *n*(1) = 0. Then while *n*(*t*) *< N* :
  a. Choose parent 1 as the best *p*_1_ = *s*(*N − t* + 1) and parent 2 as the next best *p*_2_ = *s*(*N − t*) remaining in the stack;
  b. Add *n*_*p*1, *p*2_ ∼ *ℱ*_*n*_ offspring individuals to the next generation via meiosis from their parents (*p*_1_, *p*_2_). Set *n*(*t* + 1) = *n*(*t*) + *n*_*p*1, *p*2_.
  c. Move to the next pair in the list, i.e. *t ← t* + 2.

#### 4.10.2 Compliance with Breeding recommendations

There are many reasons that breeders can be unable to comply with breeding recommendations made purely on quantitative features. These include practicalities of the geographic locations of the animals, properties of the captive individuals such as mate incompatibility, and external knowledge of suitability not encoded in the breeding model. To simulate these effects we introduce a compliance rate *c* when performing the ranking in Section 4.10.1. In stage 4, when performing pairing, with probability *c* we follow the recommendation, and otherwise (with probability 1 *− c*) we instead choose an unpaired individual (lower ranked) uniformly at random, and skip the current individual in the ranking. We examined *c* = 0.6 and *c* = 0.8.

Because fit individuals fail to mate with probability *c*, this is a strong penalty. In a real breeding program, non-compliance with a *specific mating pair* might be common (and may be 60% or lower), but the breeders are likely to find alternative good solutions that still involve that individual mating with another fit individual. Therefore we believe that this model is likely to be representative for larger *c*.

#### 4.10.3 Threshold Score Breeding Rule

A simpler process is to rank all individuals according to the selected score in the current generation, select all individuals above some threshold and then perform random mating with the selected individuals. We use a threshold value as the 50% quantile as a sensitivity analysis to the details of the breeding procedure, and the results are qualitatively the same; see Figure S2.

### 4.11 Scottish Wildcat data

The sequencing data from [20] were used by [19] where the data were analysed with MOSAIC [17] to infer the ‘local ancestry’ i.e. the population source (wildcat or domestic cat, as determined by out-group individuals and historical wildcat data) for every SNP for each individual. Briefly, the pipeline used Bowtie2 [49] for alignment to the reference (checking that the alignment rates are not affected by wildcat/domestic/hybrid status), and variants were called and filtered with GATK [50]. Relatives were removed from the reference (kinship*>* 0.125). Phasing was performed with Beagle v5.2 [14] and variants were then thinned at random to one SNP per 2kb to reduce the computational burden. We selected chromosome E2 from this dataset, resulting in 17720 SNPs for 36 Scottish wildcat and hybrid individuals.

The expected number of offspring per pair per generation is calculated from data matched to the stud book of the breeding program [47] as follows. Firstly, the observed offspring distribution per brood in the stud book is (0.23, 0.36, 0.30, 0.10, 0.01) for (1, 2, 3, 4, 5) kittens respectively, with a mean of 2.3. 30% of individuals survive to first year and average mortality is then 10% per year thereafter, with 50% of adult females bred per year, leading to a breeding probability for ages *a* = 1, …, 7 of 0.5(1 *−* 0.3)(1 *−* 0.1)^*a−*1^ with an expected number of breeding attempts of 1.83. Multiplying by the mean brood size leads to an expected number of offspring of 4.2, which we assume to be Poisson distributed.

## Code Availability

All code used in this project is available at https://github.com/danjlawson/localsancestrybreeding.

## Acknowledgements

This work was carried out using the computational facilities of the Advanced Computing Research Centre, University of Bristol - http://www.bristol.ac.uk/acrc/.

## Supplementary Figures

**Figure S1:**
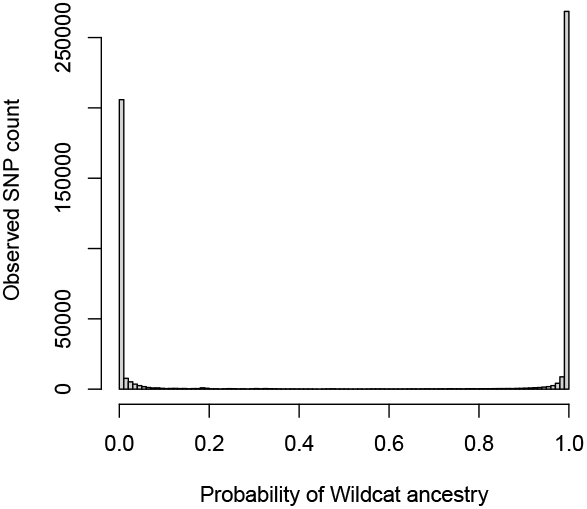
Ancestry is easy to call for hybridized species of interest. Empirical probability of wildcat ancestry (*F*_*ST*_ = 0.46) for the pre-selection Scottish wildcat data as produced by Mosaic for chromosome E5. There are no genomic positions containing considerable uncertainty (probability close to 0.5) and therefore the calls for wildcat (*p* = 1 or domestic cat (*p* = 0) are very confident and can be approximated as knowing the ancestry status. (Call accuracy is also well-calibrated as assessed in the methods paper [17]).

**Figure S2:**
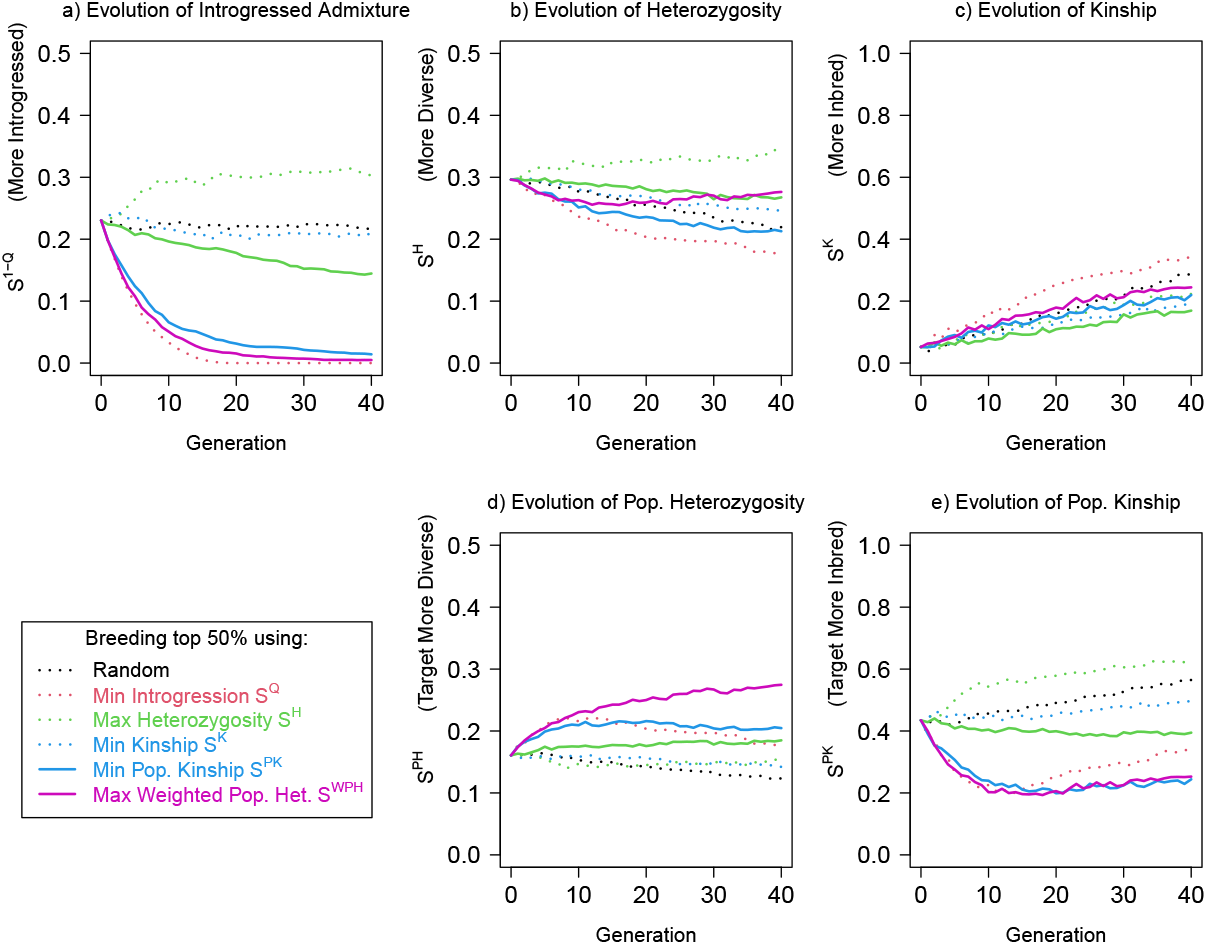
The results are robust to breeding details. Results as in Figure 3 but for a simplified breeding decision rule (Section 4.10): the top 50% of individuals by the chosen score on the *current* population are bred randomly. We quantify performance over time with scores for a) introgressed admixture fraction 1 *− S*^*Q*^, b) heterozygosity *S*^*H*^, c) kinship *S*^*K*^, d) population heterozygosity *S*^*P H*^, e) population kinship *S*^*PK*^ .

